# A lectin-based magnetic CRISPR screening protocol permits rapid discovery of genes regulating cell-surface glycosylation

**DOI:** 10.1101/2025.09.15.676433

**Authors:** Jimmy Kim, Halen Kovacs, Simon Wisnovsky

## Abstract

FACS-based CRISPR screening has emerged as a potent tool for dissecting the genetic networks that regulate cell-surface glycosylation. However, existing protocols are tedious and poorly suited to many cell models. We developed a lectin-based magnetic-activated cell sorting platform (Lec-MACS) that enables rapid identification of genes controlling expression of specific cell-surface glycans. Lec-MACS is faster and easier to perform than FACS-based screening while producing data of similar quality. We subsequently applied Lec-MACS to produce a genomic atlas of genes regulating breast cancer hypersialylation. This method will dramatically expand the scope and throughput of genetic screens targeted at cell-surface glycans.

## Introduction

All living cells are coated with carbohydrate molecules (termed *glycans*) that play crucial roles in a wide range of cellular processes^1,2^. Glycans are added to proteins and lipids on the surface of cells through the combined action of hundreds of glycosyltransferase enzymes^1-3^. In human cells, many glycans are terminated with a negatively charged, 9-carbon monosaccharide called *sialic acid*^1,2^. The addition of sialic acids onto glycan chains occurs via sialyltransferases located in the Golgi Apparatus. There also exist numerous sialidase enzymes that cleave and release sialic acids from glycoconjugates, helping to regulate surface sialylation^1,2^. Due to their position at terminal ends of glycan chains, sialic acids are heavily involved in regulating cell-cell interactions^4,5^. Sialic acid-containing glycans are thought to act as a “self-associated molecular pattern” that allow immune cells to discriminate self from non-self^4,5^.

The collection of sialylated glycans, dubbed the “sialome”, is often dysregulated in diseases such as cancer. Hypersialylation (an increase in sialic acid on the cell surface) has been observed in many carcinomas^6^. This phenomenon contributes to multiple hallmarks of cancer, such as immune evasion, mitigation of cell death, and increased tumor invasion^6,7^. For example, Siglecs (sialic acid-binding immunoglobulin-like lectins) are a family of immune receptors that have high affinity for glycans containing sialic acid^4,5^. Upon binding to sialic acid, these receptors initiate an inhibitory signaling cascade that dampens the immune response, leading to immune evasion in cancer cells^4,5^. Some studies have observed that the transcriptional upregulation of sialyltransferase genes or the downregulation of neuraminidase expression could lead to the alteration of the sialome^8^. The signaling pathways and genetic factors that coordinate these changes in cancer cells, however, are not well defined.

In previous work, we developed a genome-wide CRISPR screening method that aims to uncover potential regulators of surface sialylation in cancer cells^9^. In this approach, a cancer cell line that expresses a Cas9 construct is transduced with a genome-wide library of sgRNAs. Cells are then stained with fluorescent sialic acid-binding proteins and sorted to isolate cells exhibiting a reduction in sialylation. Subsequent sequencing of sgRNAs enriched in this low-staining population then reveals specific genes whose knockout reduces cell-surface sialylation. We have successfully applied this platform to identify genetic drivers of sialylation in hematopoietic cancer cell lines and validate new targets for cancer immunotherapy^9-13^. A similar approach has been broadly adopted by other groups to study other glycan-driven cell-surface interactions^12,14-20^.

FACS-based CRISPR screening has become a popular technique for studying glycobiology for several reasons. Firstly, a wide variety of recombinant lectins (glycan-binding proteins) can be fluorescently labeled and used to quantify glycan expression, making this approach highly modular^12,13^. Secondly, glycan biosynthesis is a tremendously genetically complex cellular phenotype. Dozens of genes encoding biosynthetic enzymes, cell-surface scaffolding genes and/or metabolic enzymes may play roles in modulating expression of a specific cell-surface glycan^12,13^. High-throughput genetic screening can interrogate thousands of genes in a single experiment, thus offering unique advantages for decoding these polygenetic circuits^12,13^. Given the large number of functional lectins encoded in both mammalian and bacterial genomes, it is likely that such techniques will become an ever-more important tool for glycobiology researchers in the future.

However, this “first-generation” approach suffers from several important drawbacks. Firstly, the FACS-based method of sorting cells is time-consuming and difficult to coordinate for large library sizes. Secondly, this method is not well-suited to the wide array of cancer cell lines that are difficult to sort in large quantities. Many adherent cancer lines exhibit significant loss of viability and poor sorting efficiency during FACS, making it difficult to perform large-scale sorting experiments. Thus, there is an urgent need for a faster, better screening method that is applicable to a wider range of cell models.

An alternative method for selecting cells from pooled CRISPR libraries is magnetic-activated cell sorting (MACS). A CRISPR screen using magnetic separation typically involves introducing a pooled library of guide RNAs into cells and labeling specific populations of interest using a specific reagent coupled to magnetic beads. A magnetic field is then applied to isolate target cells enriched or depleted for genetic perturbations that affect marker expression or cell behavior. This general strategy has been used to conduct CRISPR screens to investigate a variety of biological processes^21-23^. MACS has also been successfully applied to identify novel receptors for recombinant ligands^24^ and to map genes that regulate expression of cell-surface antigens^15,23^. However, these methods have mainly used high-affinity, protein-binding reagents like antibodies to capture cells from pooled CRISPR libraries^21-23^. It is unclear whether MACS represents a feasible strategy for studying cell surface glycosylation, where recombinant lectins are more commonly used to interrogate expression of specific carbohydrates.

In this study, we developed a rigorous protocol for optimization and implementation of MACS-based CRISPR screens using lectins (Lec-MACS). We carefully optimized key experimental parameters by performing magnetic isolations of cells with defined genetic perturbations in key glycan biosynthesis genes. This approach provides a general roadmap for researchers interested in optimizing MACS-based screening with different glycan-binding reagents. Then, we performed parallel FACS and MACS-based genetic screens in an adherent breast cancer cell line (MDA-MB-231 cells). Systematic comparison of these methods revealed that MACS produces data of similar quality. Finally, we performed a genome-wide CRISPR screen to broadly map genetic drivers of hypersialylation in breast cancer cells, an experiment that would not have been feasible using FACS-based methods. Taken together, our results provide a valuable resource for development and execution of genetic screens targeted at cell-surface glycans.

## Results

### Optimization of a system for magnetic separation of sialylated from non-sialylated cells

We first sought to develop a pipeline for optimizing magnetic separation of differentially glycosylated cells. Biosynthesis of sialic acid-containing glycans requires the action of a gene called cytidine monophosphate N-acetylneuraminic acid synthetase (CMAS)^9^. This enzyme produces an activated sugar nucleotide (CMP-Sialic Acid) that can be added to growing glycan chains by glycosyltransferases (Fig. 1A). Genetic knockout of CMAS has been shown in previous studies to totally ablate sialic acid synthesis^9,25^. To generate a cell model lacking sialoglycan expression, we thus transduced MDA-MB-231-dCas9KRAB cells with an sgRNA targeting the CMAS gene. dCas9KRAB is a transcriptional repression domain that mediates silencing of target genes. MDA-MB-231s are adherent breast cancer cells that are difficult to sort in large numbers by FACS. We therefore felt they would be an ideal model for testing the utility of a MACS-based approach to CRISPR screening. Knockdown of CMAS was confirmed by qPCR analysis, which revealed a near-complete silencing of gene expression (Fig. 1B). We then stained MDA-MB-231 cells with SiaFind pan-specific Lectenz (Lectenz Bio, hereafter termed *Lectenz*). This reagent is an engineered lectin that binds broadly to cell-surface sialic acids^26,27^. WT and CMAS KD cells were stained with Lectenz-biotin and a fluorescently labeled streptavidin conjugate. Subsequent flow cytometry analysis showed that WT MDA-MB-231 cells exhibit strong expression of sialic acid-containing glycans (Fig. 1C). CMAS KD totally ablated Lectenz binding as expected (Fig. 1C).

**Figure 1.**
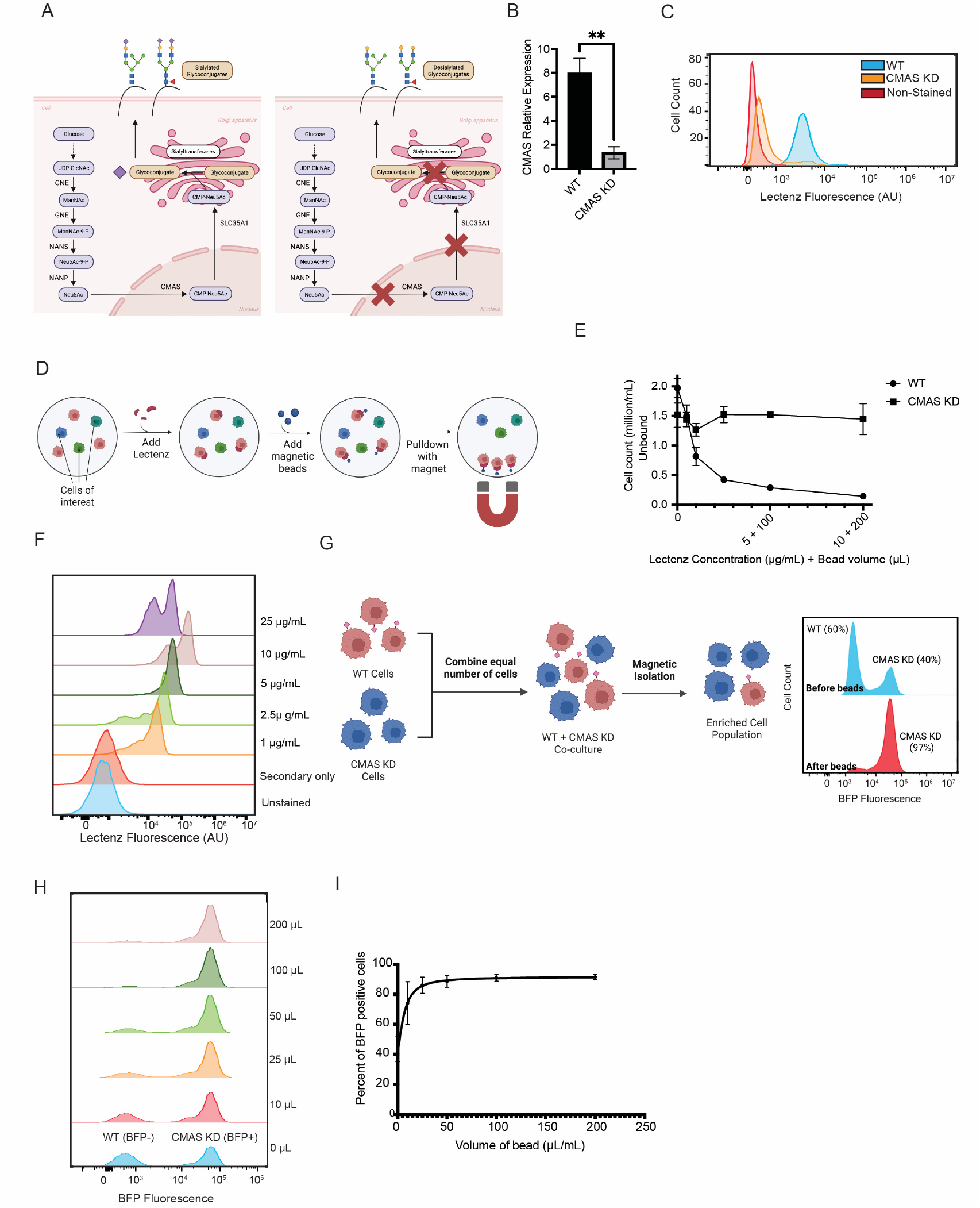
**A)** CMAS catalyzes the formation of a key intermediate in sialic acid biosynthesis. Genetic knockdown of CMAS eliminates import of sialic acid precursors into the Golgi Apparatus and prevents cell-surface sialylation. **B)** MDA-MB-231-dCas9KRAB cells were transduced with an sgRNA targeting CMAS. RNA was then extracted from these cells and levels of CMAS mRNA were quantified by RT-qPCR. **C)** MDA-MB-231-dCas9KRAB were transduced as in **B** and incubated with Lectenz for 30 minutes on ice. Cells were subsequently stained with Streptavidin DyLight488 secondary and analyzed by flow cytometry. **D)** Workflow for magnetic capture of non-sialylated cells. Cells were incubated with Lectenz as in **C** and then exposed to magnetic Streptavidin microbeads (Miltenyi). Subsequent exposure to a magnet separates cells into two populations: those which bind the beads (“Bound” fraction) and those remaining in the supernatant (“Unbound” fraction). **E)** WT and CMAS KD cells were subjected to magnetic isolation as in **D** using the indicated concentrations of Lectenz and beads. The number of cells in the unbound fraction (supernatant) was then determined using an automated cell counter. **F)** WT MDA-MB-231-dCas9KRAB cells were stained with the indicated concentrations of Lectenz as in **C**. A representative flow cytometry plot is shown **G)** WT (BFP-) and CMAS KD (BFP+) cells were mixed in a 1:1 ratio. Cells were then subjected to magnetic isolation as in **D**. A representative flow cytometry plot shows the relative proportions of WT and CMAS KD cells following isolation with the indicated cell:bead ratios. **H)** WT (BFP-) and CMAS KD (BFP+) were subjected to magnetic isolation as in **G**. A representative flow cytometry plot is provided for the indicated range of concentrations. **I)** The percentage of BFP+ cells in the unbound fraction is plotted for the indicated cell:bead ratios. All graphs show mean values plotted for n=3 biological replicates. Error bars indicate SEM.

We then explored whether magnetic isolation could be used to separate highly sialylated (WT) from lowly-sialylated (CMAS KD) cells. We labeled both WT and CMAS KD cells with Lectenz at a variety of concentrations. We subsequently incubated with Streptavidin Microbeads (Miltenyi) and separated cells into bound and unbound fractions (Fig. 1D). In WT cells, increasing the concentration of both beads and Lectenz led to a dose-dependent decrease in the number of cells remaining following isolation (the “unbound fraction”) (Fig. 1E). CMAS KD cells, conversely, were consistently retained in the unbound fraction at high densities (Fig. 1E). These results confirm that magnetic separation can be used to discriminate between lowly sialylated and highly sialylated cells. Crucially, we did find that efficient capture was highly dependent on both the concentration of Lectenz and the volume of microbeads used in the assay. At lower bead/Lectenz concentrations (2.5 µg/mL and 50 µL beads), we found that a significant number of WT cells were not captured by the beads. Higher concentrations (10 µg/mL and 200 µL beads) were required to mediate full, efficient capture of sialylated cells. This experimental design consideration is likely to be critical for effective execution of a MACS-based lectin screen.

We then determined the limiting amount of each reagent that is required for effective magnetic capture. Previous studies using magnetic isolation have found that high antigen density on the cell surface is essential for efficient MACS^15^. We therefore performed a titration experiment where cells were stained with escalating concentrations of biotinylated Lectenz. Cells were subsequently incubated with a Streptavidin-DyLight488 conjugate to allow detection of Lectenz binding by flow cytometry. Lectenz binding increased in a concentration-dependent manner up to a staining concentration of 10 µg/mL (Fig. 1F). Decreased binding and reduced cell viability was observed at higher concentrations. Lectins have been reported to induce functional changes in cell signaling in other contexts^28,29^. This effect could thus be mediated by cross-linking of adjacent sialic acid residues on the cell surface, leading to changes in receptor signaling. Regardless of the mechanism, such effects would be undesirable when performing a genomic screen, where high cell viability is required to maintain library coverage. We therefore chose to perform future experiments at a binding concentration of 10 µg/mL Lectenz.

Finally, we developed a co-culture assay to determine the concentration of beads required for efficient depletion of sialylated cells. WT and CMAS KD cells were mixed in a 1:1 ratio and stained with 10 µg/mL Lectenz. Cells were then treated with increasing volumes of Streptavidin Microbeads. The unbound fraction was isolated and analyzed by flow cytometry (Fig. 1G). As our sgRNA transfer plasmid contains a blue fluorescent protein (BFP) reporter, CMAS KD cells can be easily identified as BFP+. WT cells, conversely, displayed no BFP fluorescence. We observed significant enrichment of CMAS KD (BFP+) cells in the unbound fraction even at low bead:cell ratios (10 µL/1x10^6^ cells). Total depletion of WT cells was observed at ratios of 50 µL/1x10^6^ cells and above (Fig. 1H-I). Taken together, these data confirm that lectin-based MACS (Lec-MACS) can be used to easily separate populations of cells exhibiting differences in sialylation from a mixed cell pool.

### Magnetic and flow cytometry-based druggable genome screens map regulators of cell-surface sialylation

Having optimized our method for magnetic separation, we then performed parallel CRISPRi screens to benchmark this technique against a traditional FACS-based approach. One of the primary limitations of FACS-based screening is the maintenance of library coverage. Ideally, sorting 500-1000 cells/sgRNA is recommended for optimal data quality in a FACS-based screening experiment. When using a genome-wide library, this constraint necessitates sorting at least 100x10^6^ viable cells in a single experiment (1000 x 5 sgRNA/gene x 20,000 genes). However, cell sorters can only process a fixed number of events per second. This creates a problem for sorting genome-wide libraries. Extended sorting times make it difficult to maintain viability of cells both pre- and post-sort, as many adherent cell lines rely on anchorage-dependent signals to maintain cell health. In our hands, we have also found cell loss can occur through sorter clogging and loss of doublet cells through cell clumping.

We therefore worried that starting at a genome-wide scale would make a FACS-based screen logistically difficult to complete, preventing a head-to-head comparison with MACS. To make a meaningful comparison between FACS and MACS, we therefore chose to screen using a previously described sub-genomic library targeting ∼12,500 “druggable” genes (kinases, phosphatases, etc.)^30^. This smaller library requires sorting of many fewer cells to maintain coverage, making a FACS-based protocol easier to perform. The library also contains sgRNAs against CMAS and GNE, two sialic acid biosynthesis genes that we would expect to recover as positive control hits in both screens (Fig. 1A).

MDA-MB-231-dCas9KRAB cells were subsequently transduced with this sub-genomic library. For the flow cytometry-based screen, cells were stained with the fluorescent Lectenz reagent and sorted by FACS into two bins corresponding to the top 20% (high) and the bottom 20% (low) of the fluorescence distribution. This represents the “gold-standard” approach to lectin-based FACS screening that we and others have applied in other studies^9,11^. For the MACS-based screen, cells were harvested and subjected to magnetic separation using the basic protocol described above. A full account of our MACS separation method is given in the *Material & Methods* section. MACS separation yielded a bound fraction that adhered to the beads and an unbound fraction that remained in the supernatant. Genomic DNA was extracted from all these fractions and CRISPR libraries were amplified by PCR. Next-generation sequencing was then performed to map the abundance of sgRNAs within each distinct cell population. Parallel biological replicates were performed to track the consistency of hit recovery across different replicates.

Hit genes were identified using the Cas9 High-Throughput Maximum Likelihood Estimator (CasTLE), as described in previous studies^31^. CasTLE was chosen as it has already been applied in several high-throughput MACS-based CRISPR screens^21,32^. For the MACS screen, genes were assigned a negative effect size if sgRNAs targeting that gene were enriched in the unbound fraction, while genes were assigned a positive value if sgRNAs targeting that gene were enriched in the bound fraction. For the FACS screen, a negative effect size corresponded to sgRNA enrichment in the low fraction, while a positive effect size corresponded to enrichment in the high fraction. Knockdown of a gene with a negative effect sign thus decreased Lectenz binding. Genes were also assigned a CasTLE score, which corresponds to a measure of statistical significance for the hit gene in question. A higher CasTLE score indicates a greater level of statistical reliability. Finally, a CasTLE p-value was computed for each hit gene.

The results of both screens are presented in Figs. 2C-D and Supplementary Tables 1-2. In both screens, the top two negative regulators (those genes whose knockdown reduced sialylation) were CMAS and GNE. These are the only two genes in this sub-library that are known core components of the sialic acid biosynthesis pathway. Their recovery as hits is thus strong validation that Lec-MACS can correctly identify known regulators of glycosylation. Gene ontology (GO) analysis of top screen hits showed a similar enrichment of genes associated with processes related to glycosylation (e.g, N-acetylneuraminate metabolic process) (Fig. 2E-F). Comparing the results of these two screens provided some valuable insights into the relative strengths and weaknesses of MACS as an approach for phenotypic selection in lectin-based CRISPR screening. These comparisons are discussed in detail below.

**Figure 2.**
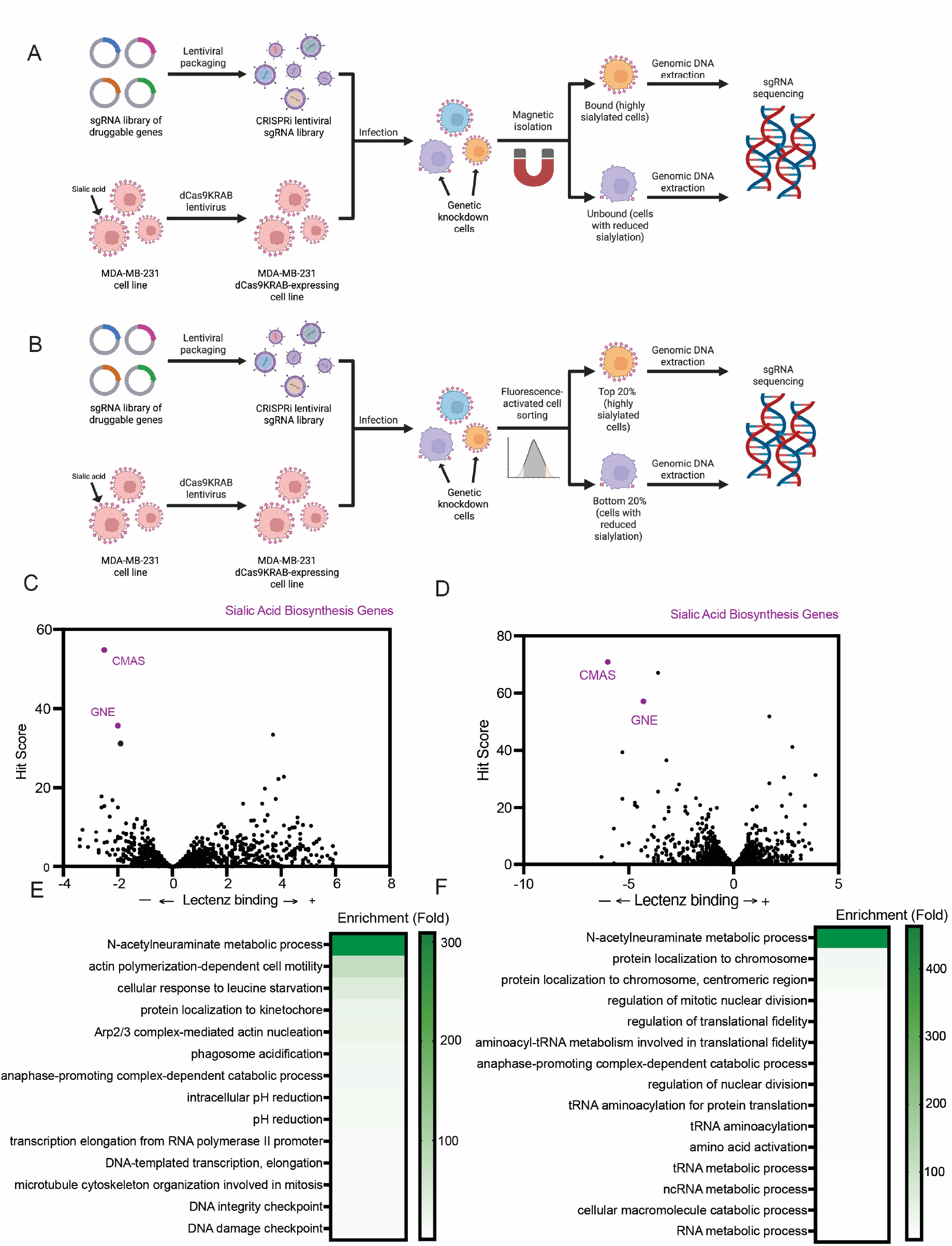
**A)** Workflow for MACS-based CRISPR screen. MDA-MB-231-dCas9KRAB cells were transduced with a targeted sub-library of sgRNAs and subjected to magnetic isolation as in **1D**. Genomic DNA was then isolated from the bound and unbound fractions. Libraries were amplified by nested PCR and subjected to next-generation sequencing. **B)** Workflow for FACS-based CRISPR screen. MDA-MB-231-dCas9KRAB cells were transduced and stained as in **1C** and sorted by FACS into high-binding (top 20% of the fluorescence distribution) and low-binding (bottom 20% of the fluorescence distribution) fractions. Genomic DNA extraction and sequencing was performed as in **2A. C)** The results of the MACS-based CRISPR screen are plotted. Each point represents a gene in the library. A higher CasTLE score (y-axis) indicates greater statistical confidence in the given hit gene. A negative effect size indicates that knockdown reduces sialylation, while a positive effect size denotes an increase in sialylation. **D)** The results of the FACS-based CRISPR screen are plotted as in **2C. E)** Enriched GO terms associated with top hits were determined for the FACS screen using GOrilla^36^. The top 15 terms with a q-value (FDR) under 0.01 are ranked by fold enrichment. **F)** Enriched GO terms associated with top hits were determined for the MACS screen as in **2E**.

### A systematic comparison of FACS & MACS-based screening

We first compared the read count distributions of samples in our screens. Fractions isolated by FACS (“Low” and “High”) and MACS (“Unbound” and “Bound”) displayed very similar read count distributions. The Gini coefficient, which measures the variation between low-count and high-count sgRNAs, was approximately 0.35 for both the MACS and FACS samples (Fig. 3A-B). This value is in the typical normal range for previously reported CRISPR screens utilizing growth-based phenotypic selection^33^. sgRNA counts were similarly distributed in each replicate, with no skewing towards outlier sgRNA counts in any sample (Fig. 3C-D). These data indicate that both our FACS and MACS protocols can maintain comparable sgRNA representation and produce similar levels of sgRNA enrichment following selection.

**Figure 3.**
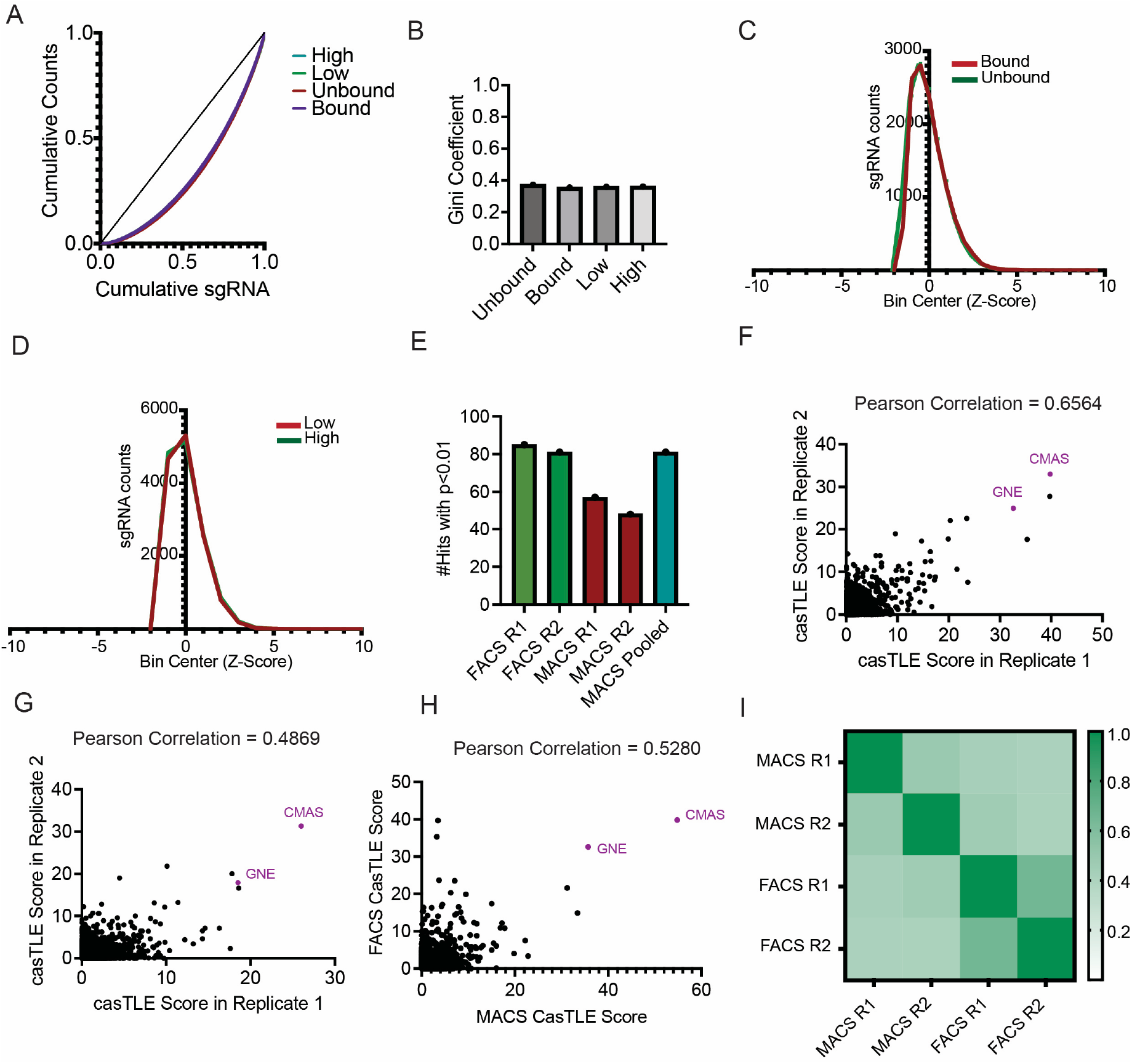
**A)** The Lorenz curve for is plotted for each isolated fraction from our MACS and FACS-based screens. The linear correlation line represents the curve for a theoretical distribution where all sgRNA counts are completely equal. **B)** The Gini Coefficients for the MACS and FACS-based screens were calculated using the Lorenz curves displayed in 3A. **C)** Normalized read count distributions for the MACS-based screen are plotted. **D)** Normalized read count distributions for the FACS-based screen are plotted. **E)** The number of hits with a CasTLE p-value < 0.01 is plotted for replicates 1 & 2 of the FACS screen (FACS R1/R2), replicates 1 & 2 of the MACS screen (MACS R1/R2), and the combined analysis of both MACS screen replicates (MACS pooled). **F)** CasTLE scores are plotted for every gene in each replicate of the FACS screen. The x-axis corresponds to R1, the y-axis corresponds to R2. **G)** CasTLE scores are plotted for every gene in each replicate of the MACS screen. The x-axis corresponds to R1, the y-axis corresponds to R2. **H)** The combined CasTLE scores are plotted for every gene in the MACS screen (x-axis) and the FACS screen (y-axis). **I)** The heat map indicates the Pearson Correlation Coefficients between each of the screen replicates included in this study.

We next compared rates of hit recovery between the two screening approaches. We started by measuring the number of statistically significant hits in each screen. Both individual replicates of the FACS screen produced a greater number of hits (CasTLE p value <0.01) than either individual replicate of the MACS screen (Fig. 3E). This result was not surprising, as the FACS method of selection intrinsically allows more stringent separation of cell populations than is possible with MACS. However, pooled analysis of two MACS screen replicates produced rates of hit recovery that were greater than either individual replicate of the FACS screen. Here, we would emphasize that performing multiple replicates of a MACS screen in parallel is relatively trivial. Adding additional replicates of a FACS screen, conversely, significantly increases experimental complexity and sort time. In our hands, performing two parallel replicates of MACS-based selection still consumed significantly less time than a single round of FACS-based selection. Lec-MACS can thus identify hit genes at comparable rates to FACS-based screening on a time-normalized basis.

Finally, we tested the reproducibility of these screening approaches across multiple replicates. Here, we noted a common limitation of both FACS and MACS-based screening methods. For both our FACS and MACS screens, CasTLE scores in replicates 1 and 2 were only moderately correlated (PCC=0.65 and 0.49 respectively) (Fig. 3F-G). When compared against each other, the FACS screen and the MACS screen displayed a similar level of correlation (0.52) (Fig. 3H-I). CMAS and GNE were among the top hits in all our replicates, indicating that both approaches can identify and rank the strongest expected hits with good reproducibility. However, there was considerable intra-replicate variability in the relative ranking of genes outside of this top tier. Both FACS and MACS-based screening are thus associated with significant statistical noise, especially at lower hit scores. This factor must be considered when selecting hits for downstream validation.

These PCC values are notably lower than those typically reported with drug toxicity or gene essentiality screens, where intra-replicate correlations of 0.8 or above are often obtained^33^. In these screens, however, it is much more common to perform multiple rounds of selection over longer times, which improves the signal to noise ratio^33^. Performing multiple, sequential selections is quite difficult when using FACS, as it would be challenging to re-seed and expand enough viable cells after sorting without bottlenecking a large library. While we did not explore doing multiple MACS-based selections in this study, this approach has been successfully adapted in other MACS-based screens^24,32^. The ability to easily perform multiple, sequential selections to improve data quality is yet another clear advantage of MACS over FACS-based selection. Whether this step is necessary will vary according to user preference and study design, but the option clearly exists for future users of Lec-MACS.

These results validate Lec-MACS as an alternative strategy for probing genetic regulators of glycosylation. The output and quality of data produced by Lec-MACS are similar to traditional FACS in most respects. Based on these considerations, we would recommend Lec-MACS as a superior protocol for most applications. However, we would urge users of Lec-MACS to complete multiple parallel screens and to analyze data using a software like MaGeCK or CasTLE that permits pooled statistical analysis of multiple screen replicates^31,33^. We would also suggest focusing validation on hits that produced strong effects in multiple biological replicates. In our view, this approach represents the ideal compromise between speed, ease of use and data quality.

### A genome-wide Lec-MACS screen maps the genetic landscape of sialylation in an adherent cancer cell line

Having extensively optimized and benchmarked our new Lec-MACS protocol, we finally wanted to demonstrate its utility for large-scale genomic screening studies. As mentioned above, conducting FACS-based genome-wide CRISPR screens in adherent cancer cell lines is technically challenging. In our hands, many cell lines and primary cell adherent cells simply cannot be stained with fluorescent lectins and sorted effectively by FACS while maintaining adequate coverage for a genome-wide screen. Lec-MACS, conversely, can be carried out quickly using simple benchtop equipment. The time needed to conduct magnetic isolation is also insensitive to the size of the library. Lec-MACS can thus be easily scaled to quite large genome-wide screening experiments. The capacity to conduct genome-wide glyco-genomic screens in adherent cell lines would thus represent a clear advance for Lec-MACS over prior methods.

We therefore conducted a genome-wide screen to map genetic regulators of cell-surface sialylation across the whole genome. To our knowledge, this is the first such screen to be conducted in a non-hematopoietic cell model. MDA-MB-231-dCas9KRAB cells were transduced with a genome-wide CRISPRi library (104,535 sgRNAs, 5 sgRNAs/gene). After selection and expansion, cells were then subjected to the Lec-MACS pipeline that has been described extensively above. 2x10^8^ cells were processed by MACS, corresponding to approximately 2000x coverage relative to the size of the pooled library. All steps of the cell isolation protocol (e.g, cell harvesting, incubation with Lectenz/beads and DNA extraction) were completed in approximately 5 hours. Following nested PCR and sequencing, screen results were analyzed via CasTLE as described above (Fig. 4A, Supplementary Table 3).

**Figure 4.**
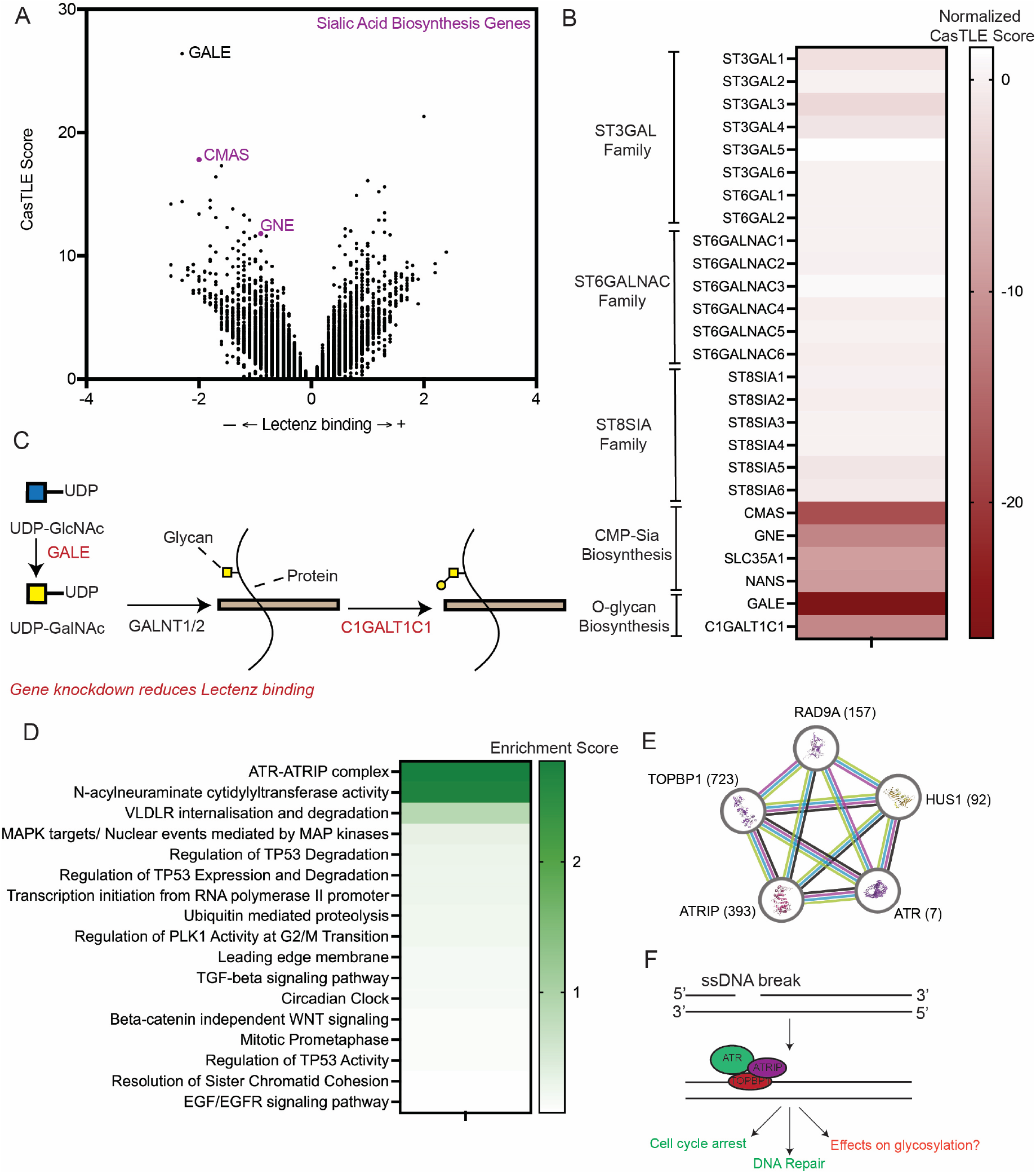
**A)** The results of the MACS-based genome-wide CRISPR screen are plotted. Each point represents a gene in the library. A higher CasTLE score (y-axis) indicates greater statistical confidence in the given hit gene. A negative effect size indicates that knockdown reduces sialylation, while a positive effect size denotes an increase in sialylation. **B)** CasTLE scores for all known sialyltransferase genes in the human genome, as well as known CMP-Sialic Acid and O-glycan biosynthesis genes. Scores have been given a sign to indicate directionality of effect. A negative score indicates that gene knockdown decreased Lectenz binding. **C)** Biosynthetic pathway for generation of UDP-GalNAc intermediates and O-linked glycosylation of cell-surface glycoproteins. The roles of GALE and C1GALT1C1 in this process are clearly indicated. **D)** Functional terms associated with top negative hits were determined using the STRING database. The enrichment score quantifies how much the values associated with a specific term deviate from the dataset’s average. All terms listed were statistically significant with an FDR < 0.01. **E)** Network analysis showing characterized interactions between known components of the ATR-ATRIP complex (GOCC:0070310). The ranking of each gene hit in our CRISPR screen is indicated. **F)** The biological function of the ATR-ATRIP complex in ssDNA break repair is indicated.

Overall, our genome-wide screen identified 331 significant hits (P<0.01). Once again, CMAS and GNE were two of the top negative regulators identified in our dataset. In addition, we also noted the presence of several other key sialic acid biosynthesis genes that had not been represented in our targeted sub-library. These included SLC35A1, which is the nucleotide sugar transporter that mediates import of CMP-sialic acid into the Golgi, and NANS, which functions in sialic acid biosynthesis upstream of CMAS (Fig. 1A). Taken together, these data strongly indicate that Lec-MACS can reveal known regulators of cell-surface sialylation on a genome-wide scale.

In contrast, we did not identify any sialyltransferase (ST) enzymes as significant hits. STs play a key role in sialylating specific glycan chains and glycoproteins (Fig. 4B). However, there are over 20 of these genes in the human genome. Many STs also act on overlapping substrates. This means that genetic inhibition of any one ST may produce muted effects on cell-surface sialylation if other redundant STs are co-expressed in the same cell. The Lectenz reagent we used for our screens has also been shown to have broad binding specificity to many underlying sialoglycan structures^26,27^. The absence of ST enzymes as top hits strongly implies that TNBC cells remodel their cell-surface glycome through fundamental changes in monosaccharide metabolism and biosynthesis, rather than by transcriptional upregulation of any individual ST.

Further supporting this point, our genome-wide screen identified several core genes involved in O-linked glycan biosynthesis (Fig. 4B-C). GALE, for example, is a key epimerase that converts UDP-GlcNAc to UDP-GalNAc, a unique building block in the biosynthesis of O-linked glycans. C1GALTC1, additionally, is a critical enzyme involved in extension and branching of O-linked glycan structures. We did not identify any genes of comparable importance to biosynthesis of N-linked glycans or glycolipids. These results indicate that cell-surface sialic acids in TNBC cells may be selectively enriched at the terminal end of O-linked glycan chains. If so, this finding could have important implications for understanding how sialic acids function to promote breast cancer tumorigenesis.

Finally, we sought to identify broader biological processes that may play a novel role in regulation of sialoglycan biosynthesis in breast cancer cells. Using the STRING database, we performed functional enrichment analysis on our ranked list of gene hits. We focused our analysis on functional terms that were selectively enriched among hits with a negative effect score (those whose knockdown reduced cell-surface sialylation). Terms related to O-glycan and sialic acid biosynthesis were highly enriched, as expected (Fig. 4D). This analysis also revealed an unexpected result: the distinct enrichment of gene hits that are known components of the ATR-ATRIP complex. This protein complex plays a key role in DNA damage repair and the protection of cells from replication stress. Six known components of this complex (ATR, ATRIP, TOPBP1, HUS1 and RAD9A) all showed significant negative effect sizes in our screen (Fig. 4E-F). ATR, a kinase whose activity is essential for DNA damage response signaling, was amongst the top hits of our screen. While not confirmed, this result implies an interesting novel link between the DNA damage response and regulation of cell-surface sialylation.

Taken together, all our results conclusively prove that Lec-MACS can rapidly characterize both known and unknown regulatory pathways that modify cell-surface glycosylation patterns. These data validate Lec-MACS as a robust tool for genome-wide screening and target discovery in adherent cancer cell lines.

## Discussion

Our studies reveal several considerations for glycobiology researchers choosing between FACS- and MACS-based screening approaches. Indeed, ours is one of the only studies to rigorously compare these two methods using the same cell surface-binding reagent. FACS does offer better rates of hit discovery on a per-replicate basis. However, Lec-MACS was still able to identify known positive control genes as well as several novel hits of interest with comparable efficiency to FACS. Lec-MACS is also clearly superior in basic parameters of speed, versatility and multiplexing capacity. This aspect makes it relatively straightforward to improve data quality by performing additional replicates. The choice of protocol will thus depend on the biological context and experimental model. FACS-based screening will still represent the ideal option when working with small sgRNA libraries and/or cell line models that are easily amenable to FACS. However, we view MACS-based screening as a superior technique for most applications, such as when working with adherent cancer cell lines, primary cells, and/or large genome-wide CRISPR libraries.

Our results also emphasize that careful optimization of a magnetic separation protocol is critical for success. Before embarking on a magnetic selection-based CRISPR screening study, it is essential to develop a pilot system for optimizing efficient capture of genetically modified cells. This allows screens to be conducted in the confidence that MACS will effectively enrich cells with a known phenotype of interest. Our approach provides a general strategy for developing such protocols using a diverse array of lectins and glycan-binding antibodies. Taken together, our results thus provide a validated, generally applicable strategy for optimizing and conducting genetic screens using recombinant glycan-binding reagents.

Additionally, there are some obvious modifications to the MACS protocol that may further improve its reliability. Above, we have already discussed some of the statistical disadvantages that are produced by performing single rounds of selection with FACS and MACS. For MACS-based screening, this problem could be mitigated simply by pooling larger numbers of biological replicates. This would be relatively straightforward to do given the speed of the MACS-based separation protocol. Alternatively, we could also use MACS to perform multiple, sequential isolation steps where cells are expanded and re-cultured in between isolations. This step may improve the reliability of hit recovery and help to enrich sgRNAs that induce more moderate phenotypes. Sorting and culturing viable cells by FACS would be extremely difficult for many cell lines, so this option may further improve the relative appeal of MACS vs. FACS-based screening. Lastly, exploring additional types of beads (nanobeads vs. microbeads vs. magnetic columns) as well as different sialic-acid binding reagents could be a further step for methods development. We therefore see Lec-MACS as a technology with significant future promise.

In this study, we focused primarily on method development, benchmarking and validation. However, our screens also revealed some notable novel findings that will be exciting to pursue and characterize in future work. Our genome-wide screen, for example, revealed a putative functional role for the ATR-ATRIP complex in regulating cell surface sialylation. We want to emphasize that this result has not been extensively validated, and all CRISPR screen results are prone to artifacts and false positives. If confirmed, however, this result could reveal fascinating novel crosstalk between stress response signaling and cell-surface glycosylation. Inhibition of ATR signaling has been shown in prior work to induce widespread DNA replication stress. Such stress can induce upregulation of ligands for activating innate immune receptors, thus facilitating detection and destruction of damaged cells^34,35^. Our results suggest that the DNA damage response may also regulate immune recognition through modulating expression of immune-inhibitory sialoglycans. While preliminary, this hypothesis will be exciting to test and validate in future work.

Finally, we envision numerous future opportunities to apply our magnetic screening platform towards discovery of druggable targets. Crucially, Lec-MACS will allow parallel genomic screens to be conducted rapidly in multiple model cell lines representing different genetic backgrounds and disease phenotypes. This advance will allow much more extensive, in-depth characterization of the “master-regulators” of cancer glycome remodeling in a range of disease models. The potential to discover new, high-value druggable targets from such studies is extremely high.

## Acknowledgements

S.W. is supported by funding from the Canadian Institutes of Health Research (CIHR), the National Sciences & Engineering Research Council of Canada (NSERC), the Canadian Cancer Society (CCS), the Cancer Research Society (CRS) and the Canadian Glycomics Network (GlycoNet)

## Materials & Methods

### Cell culturing and conditions

Human MDA-MB-231 triple negative human breast adenocarcinoma (ATCC) and Lenti-X™ HEK293T cells (Takara Bio) were cultured in Dulbecco’s modified eagle’s (DMEM) medium (Gibco, 11995065) supplemented with 10% fetal bovine serum (FBS) (Gibco). The cell lines were cultured in a humidified incubator set at 37°C and 5% CO_2_. All cell lines were tested for mycoplasma prior to experiments.

### Plasmid construction for CRISPRi

A single guide RNA (sgRNA) sequence that targeted the CMAS gene was cloned into an expression vector containing sequences encoding a puromycin resistance gene and blue fluorescent protein (BFP) (Addgene #60955) using a previously described protocol^11^. sgRNA sequences are provided below.

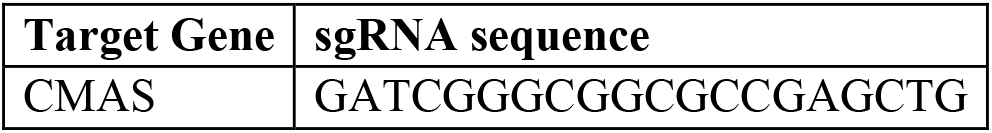

### Lentiviral transfection and transduction

For CRISPRi sublibrary virus preparation, 7.5×10^6^ HEK293T cells were plated into a 150mm plate and left to adhere to the plate overnight. A transfection mix consisting of 1.3 mL of Opti-MEM (Gibco, 31985062), 8 μg of sublibrary plasmid (Addgene, #83971), 4 μg of psPAX2 (Addgene #12260), 4 μg of pMD2.G (Addgene #12259) and 48 μL of TransIT-LT1 transfection reagent (Mirus) was assembled. The mix was added in a dropwise fashion into the culture medium of the HEK293T cells, then cells were placed back into the incubator for 72 hours. After transfection, the supernatant was collected, spun down at 3000×g for 10 minutes, then carefully collected and aliquoted into -80°C until viral transduction.

For CRISPRi genome-wide library virus preparation, 7.5×10^6^ HEK293T cells were plated into three 150mm plates and left to adhere to the plate overnight. A transfection mix consisting of 1.3 mL of Opti-MEM (Gibco, 31985062), 8 μg of the library plasmid (Addgene, #83971-83976), 4 μg of psPAX2 (Addgene #12260), 4 μg of pMD2.G (Addgene #12259) and 48 μL of TransIT-LT1 transfection reagent (Mirus) was assembled per plate. The mix was added in a dropwise fashion into the culture medium of the HEK293T cells, then cells were placed back into the incubator for 72 hours. After transfection, the supernatant was collected, spun down at 3000×g for 10 minutes, then the supernatant was concentrated 5-fold using a lentivirus concentration reagent (Takara Bio). The concentrated supernatant was then stored in -80°C until viral transduction.

For sgRNA expression and dCas9-KRAB construct virus preparation, 5×10^5^ HEK293T cells were plated into one well of a 6-well plate and left to adhere to the plate overnight. A transfection mix consisting of 500 μL of Opti-MEM, 2 μg of sgRNA-encoding plasmid, 1 μg of psPAX2 (Addgene #12260), 1 μg of pMD2.G (Addgene #12259) and 7.5 μL of TransIT-LT1 transfection reagent (Mirus) was assembled. The mix was added in a dropwise fashion into the culture medium of the HEK293T cells, then cells were placed back into the incubator for 72 hours. After transfection, the supernatant was spun at 3000×g for 10 minutes, then concentrated down to 1 mL using the Lenti-X Concentration Reagent and aliquoted into -80°C until viral transduction.

### Generation of MDA-MB-231 dCas9-KRAB-expressing cell line

Wild-type MDA-MB-231 cells were transduced with lentivirus containing a plasmid encoding an inactive Cas9 fused to a KRAB repressor domain (dCas9-KRAB) (Addgene #122205). The cell line was then selected using media supplemented with 5 μg/mL blasticidin (InvivoGen).

### Generation of CRISPR knockdown (CRISPRi) cell lines

To generate single gene knockdown (KD) cell lines, MDA-MB-231 cells stably expressing dCas9-KRAB were harvested using TrypLE. 5×10^5^ cells were transduced with the respective sgRNA-containing viral supernatant and growth media supplemented with 8 μg/mL polybrene (Sigma) in a 6-well plate for 48 hours. Before selection, BFP expression of cells was analyzed by flow cytometry to check for transduction efficiency. Cells were then selected over 96 hours in media supplemented with 1μg/mL puromycin (InvivoGen).

### Generation of CRISPRi sublibrary cell line

Before generating the sublibrary-containing cells, the virus containing the sublibrary was titered to calculate the volume of lentiviral supernatant needed to achieve 0.3 MOI. For transduction, MDA-MB-231 cells stably expressing dCas9-KRAB were harvested using TrypLE. To meet 1000x coverage of the guides encoded in the library, 3×10^7^ cells were transduced with appropriate volume of titered viral supernatant and growth media supplemented with 8 μg/mL polybrene (Sigma) across four 150mm plates for 48 hours. BFP expression of cells was analyzed by flow cytometry for transduction efficiency. Cells were then selected over 96 hours in media supplemented with 1μg/mL puromycin (InvivoGen).

### qPCR analysis

Primers were designed to amplify the protein coding regions of the gene of interest. Cells to be analyzed were pelleted and washed with PBS. The total RNA was extracted from each sample (Monarch® Total RNA Miniprep Kit, NEB, T2010S) followed by cDNA synthesis (LunaScript® RT SuperMix Kit, NEB, E3010S) using protocols supplied by the manufacturer. Reactions were set up with a ready-to-use qPCR master mix (Luna® Universal qPCR Master Mix, NEB, M3003S). The qPCR reactions were run for two hours (StepOnePlus Real-Time PCR System, Applied Biosystems) then the resulting Ct values were analyzed using GAPDH as a housekeeping gene. Primers are provided in the table below.

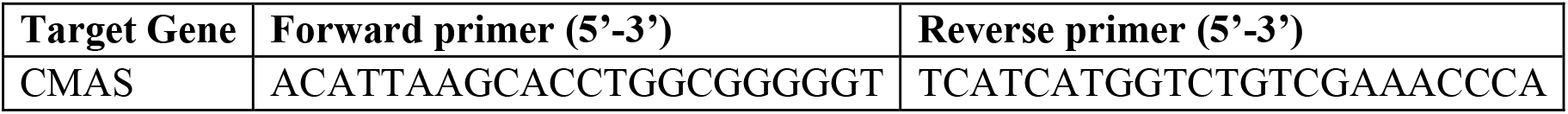

### Detection of cell surface sialylation using SiaFind-Lectenz

Cells were harvested with TrypLE (Thermo), washed with PBS, then resuspended at 1×10^6^ cells/mL. Cells were aliquoted in 100 µL into a 96-well V-bottom plate, then centrifuged at 600×g for 5 minutes, aspirated, then resuspended in 100 μL of 10 μg/mL SiaFind Pan-Lectenz (Lectenz Bio) and incubated on ice for 30 minutes with gentle mixing every 10 minutes. After the incubation, cells were pelleted, then resuspended in 100 μL of diluted DyLight™ 488 Streptavidin (DyLight 405218) (1:1000) and incubated on ice for 30 minutes with gentle inversion every 10 minutes. Following the incubation, cells were washed twice with PBS then analyzed using a CytoFLEX LX flow cytometer to measure the median fluorescence intensity of FITC.

### Magnetic separation of cells

Wild-type MDA-MB-231 dCas9-KRAB cells and CMAS KD MDA-MB-231 cells were harvested using TrypLE and washed twice with PBS before resuspending to 5×10^6^ cells/mL. 2.5×10^6^ cells of each population were combined and aliquoted into 1.5 mL eppendorf tubes. The co-culture was spun down and resuspended in PBS containing SiaFind Pan-Lectenz (Lectenz Bio) and incubated on ice for 30 minutes with gentle inversion every 10 minutes. After the incubation, cells were pelleted and resuspended in PBS containing the indicated volume Streptavidin Magnetic Beads (Miltenyi) at a range of bead concentrations. The cells were vortexed to mix and incubated for 15 minutes on a rotating platform in a cold room. After incubation, the culture tube containing magnet-labeled cells was placed inside an EasySep Magnet (StemCell) for two minutes, then the supernatant was carefully transferred (without touching the sides of the tube) to a clean culture tube. The new tube was placed in the magnet again for two minutes, then the supernatant was collected in a 15 mL Falcon tube (unbound sample). The initial tube, now containing cells bound to the sides of the tube, was thoroughly rinsed with PBS to dislodge cells and set aside (bound sample). The unbound fraction was collected and analyzed through the CytoFLEX LX flow cytometer to calculate the percentage of BFP-positive cells (CMAS KD) in the unbound fraction.

### CRISPRi sublibrary screening through MACS

Each screen was carried out in technical duplicates. 6×10^7^ MDA-MB-231 H1 sublibrary transduced cells were harvested with TrypLE (Thermo), washed with PBS, then resuspended at 5×10^6^ cells/mL in 12 mL PBS. 6 mL of the cell suspension (3×10^7^ cells) was aliquoted into two 15 mL Falcon tubes. Cells were centrifuged at 600×g for 5 minutes, aspirated, then resuspended in 6 mL of 10 μg/mL SiaFind Pan-Lectenz (Lectenz Bio) and incubated on a rocker in a cold room for 30 minutes. After the incubation, cells were pelleted then resuspended in 6 mL solution consisting of 4.8 mL PBS and 1.2 mL Streptavidin Magnetic Beads (Miltenyi), then vortexed to mix. Cells were strained through a 70μm filter to remove any clumps prior to sorting. The cell-bead suspension was divided into two 5 mL culture tubes (3 mL each) then incubated for 30 minutes on a shaker in a cold room. After incubation each tube was subjected to magnetic separation as described above. The genomic DNA of the unbound and bound samples were immediately extracted using the PureLink™ Genomic DNA Mini Kit (Invitrogen) following the manufacturer’s protocol. Unbound and bound samples were pooled following extraction to yield a single Unbound fraction and a single Bound fraction.

### CRISPRi sublibrary screening through FACS

Each screen was carried out in technical duplicates. 6×10^7^ MDA-MB-231 H1 sublibrary transduced cells were harvested with TrypLE (Thermo), washed with PBS, then resuspended at 5×10^6^ cells/mL in 12 mL PBS. 6 mL of the cell suspension (3×10^7^ cells) was aliquoted into two 15 mL Falcon tubes. Cells were centrifuged at 600×g for 5 minutes, aspirated, then resuspended in 6 mL of 10 μg/mL SiaFind Pan-Lectenz (Lectenz Bio) and incubated on a rocker in a cold room for 30 minutes. After the incubation, cells were pelleted, then resuspended in 6 mL of diluted DyLight™ 488 Streptavidin solution (1:1000) and incubated on a rocker in a cold room for 30 minutes. The cells were pelleted and washed twice, then resuspended in 6 mL PBS. Cells were strained through a 70 μm filter to remove any clumps prior to sorting. The cells were sorted using the Astrios sorter (Beckman Coulter) and gating strategy was as follows: gate for live cells, then single cells, then BFP positive cells, then top and bottom 20% of cells were sorted into respective 15 mL Falcon tubes containing 2 mL DMEM supplemented with 10% FBS. Genomic DNA was immediately extracted following the sort using the PureLink™ Genomic DNA Mini Kit (Invitrogen) following the manufacturer’s protocol.

### CRISPRi genome-wide MACS screening

2×10^8^ MDA-MB-231 CRISPRi genome-wide library transduced cells were harvested with TrypLE (Thermo), washed with PBS, then resuspended at 5×10^6^ cells/mL in 40 mL PBS. Cells were centrifuged at 600×g for 5 minutes, aspirated, then resuspended in 40 mL of PBS containing 10μg/mL SiaFind Pan-Lectenz (Lectenz Bio) and incubated on a rocker in a cold room for 30 minutes. After the incubation, cells were pelleted then resuspended in 40 mL solution consisting of 36 mL PBS and 4 mL Streptavidin Magnetic Beads (Miltenyi), then vortexed to mix. Cells were strained through a 70 μm filter to remove any clumps prior to sorting. The cell-bead suspension was then divided across 14 5 mL culture tubes (3 mL each) then incubated for 30 minutes on a rotating platform in a cold room. After incubation each tube was subjected to magnetic separation as described above, then the unbound and bound samples from each tube were pooled together. The genomic DNA of the unbound and bound samples were immediately extracted using the QIAamp DNA Blood Maxi Kit (Qiagen) following the manufacturer’s protocol.

### Library amplification, sequencing and data processing

The guide sequences for each population of cells were amplified and barcoded through a nested PCR using a previously described protocol^11^. Primers for both outer and inner PCRs are given below. PCR amplicons were gel purified, then Custom Sequencing was performed on either an Illumina NextSeq 550 platform or Oxford Nanopore Technology (Plasmidsaurus) with custom analysis and annotation. Libraries were sequenced at a coverage greater than 200 reads/sgRNA. The resulting FASTQ files were processed using SeqKit2^37^ to filter for reads between 20 and 500 bps. A reverse complement of each FASTQ file was also generated, then both original and reverse complemented FASTQs were aligned to a library file using guide-counter (https://github.com/fulcrumgenomics/guide-counter). The sgRNA counts from each sample pair were pooled and compared using CasTLE (Cas9 High Throughput maximum Likelihood Estimator)^31^.

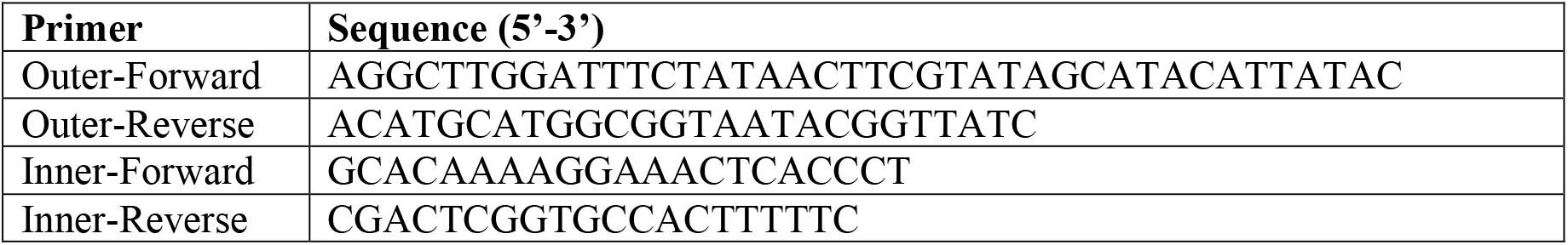

### Functional enrichment analysis

For the GO term analysis in Fig. 2, hit genes were ranked by CasTLE score and term discovery was performed using GOrilla^36^. For the broad functional enrichment analysis in Fig. 4, genes were first assigned a normalized CasTLE score, where the gene’s CasTLE score was multiplied by the sign of the effect size. The genes were then ranked with the lowest (most negative) normalized score at the top. Functional enrichment was then performed using the STRING database (https://string-db.org/).

